# In vivo proximity proteomics uncovers palmdelphin (PALMD) as a Z-line-associated mitigator of isoproterenol-induced cardiac injury

**DOI:** 10.1101/2023.12.06.570334

**Authors:** Congting Guo, Blake D. Jardin, Junsen Lin, Rachelle L. Ambroise, Ze Wang, Luzi Yang, Neil Mazumdar, Fujian Lu, Qing Ma, Yangpo Cao, Canzhao Liu, Xujie Liu, Feng Lan, Mingming Zhao, Han Xiao, Erdan Dong, William T. Pu, Yuxuan Guo

## Abstract

Z-lines are core ultrastructural organizers of cardiomyocytes that modulate many facets of cardiac pathogenesis. Yet a comprehensive proteomic atlas of Z-line-associated components remain incomplete. Here, we established an adeno-associated virus (AAV)-delivered, cardiomyocyte-specific, proximity-labeling approach to characterize the Z-line proteome in vivo. We found palmdelphin (PALMD) as a novel Z-line-associated protein in both adult murine cardiomyocytes and human pluripotent stem cell-derived cardiomyocytes. Germline and cardiomyocyte-specific *palmd* knockout mice were grossly normal at baseline but exhibited compromised cardiac hypertrophy and aggravated cardiac injury upon long-term isoproterenol treatment. By contrast, cardiomyocyte-specific PALMD overexpression was sufficient to mitigate isoproterenol-induced cardiac injury. PALMD ablation perturbed transverse tubules (T-tubules) and their association with sarcoplasmic reticulum, which formed the Z-line-associated junctional membrane complex (JMC) essential for calcium handling and cardiac function. These phenotypes were associated with disrupted localization of T-tubule markers caveolin-3 (CAV3) and junctophilin-2 (JPH2) and the reduction of nexilin (NEXN) protein, a crucial Z-line-associated protein that is essential for both Z-line and JMC structures and functions. PALMD was found to interact with NEXN and enhance its protein stability while the *Nexn* mRNA level was not affected. Together, this study discovered PALMD as a potential target for myocardial protection and highlighted in vivo proximity proteomics as a powerful approach to nominate novel players regulating cardiac pathogenesis.

**Highlights:**

- In vivo proximity proteomics uncover novel Z-line components that are undetected in in vitro proximity proteomics in cardiomyocytes.
- PALMD is a novel Z-line-associated protein that is dispensable for baseline cardiomyocyte function in vivo.
- PALMD mitigates cardiac dysfunction and myocardial injury after repeated isoproterenol insults.
- PALMD stabilizes NEXN, an essential Z-line-associated regulator of the junctional membrane complex and cardiac systolic function.

## Introduction

Myofibrils are the specialized actomyosin complexes that drive the contraction and relaxation of muscle cells. In cardiomyocytes, the structural and functional units of myofibrils are called sarcomeres, which are interconnected at their boundaries named Z-lines (Lange et al., 2006; Sanger and Sanger, 2008). Z-lines are assembled by α-actinin-2 (ACTN2), which crosslinks anti-parallel actin filaments from adjacent sarcomeres, providing anchorage sites for an array of other sarcomere-associated proteins. In addition to the structural role of Z-lines, Z-lines are a hub for signal transduction and gene expression regulation (Frank and Frey, 2011; Guo et al., 2021). Z-lines provide the scaffold for other critical subcellular structures, particularly the transverse tubule (TT) and the sarcoplasmic reticulum (SR). TTs are tubular, transverse invaginations of plasma membrane that facilitates the propagation of action potentials from the cell surface into its interior to activate the L-type calcium channel Ca_V_1.2. The SR is a specialized endoplasmic reticulum (ER) that stores calcium and mediates calcium release via the calcium channel ryanodine receptor 2 (RYR2). TT and SR membranes closely associate with each other via junctophilin-2 (JPH2) at microdomains known as the junctional membrane complexes (JMCs). JMCs facilitate calcium-induced calcium release by closely juxtaposing Cav1.2 on the TT with RYR2 on the SR, resulting in the close coupling of cardiac excitation to Ca^2+^ release that is essential for cardiomyocyte contractility (Hong and Shaw, 2017; London, 2017).

JMCs are located at subdomains of TTs located along the Z-lines, but how Z-lines interplay with JMCs to modulate cardiomyocyte function remains poorly understood. Nexilin (NEXN) likely mediates the crosstalk between Z-lines and JMCs. On one hand, NEXN is known as an actin filament-binding protein that maintains Z-line stability (Hassel et al., 2009). On the other hand, NEXN associates with JMCs and is required for TT formation (Liu et al., 2019). Human genetic mutations of *NEXN* cause either dilated or hypertrophic cardiomyopathy (Hassel et al., 2009; Wang et al., 2010). However, little is known about the regulatory mechanisms of NEXN-mediated function involving Z-lines, TTs and JMCs.

Comprehensive mapping of Z-line-associated proteins is pivotal to elucidate the molecular mechanisms underlying cardiac pathophysiology. Proximity labeling is recently emerging as a powerful approach to depict the proteomics of the Z-line and TT/JMC subdomains (Kushner et al., 2022; Rees et al., 2015). This technology relies on a promiscuous biotin ligase such as BirA* or BioID2 (Kim et al., 2016; Roux et al., 2012) or an engineer peroxidase such as APEX2 (Lam et al., 2015) to biotinylate proteins adjacent to a given bait. The labeled proteins are then purified by their binding to streptavidin, followed by identification and quantification by mass spectrometry (MS) (Kim et al., 2016; Roux et al., 2012).

To date, multiple proximity proteomics analyses have been reported to characterize the Z-line, TT or JMC-associated baits. These protein baits include Ca_V_1.2 (Liu et al., 2020), JPH2 (Feng et al., 2020), and ACTN2 (Ladha et al., 2021). One major technical concern about cardiomyocyte proximity proteomics involves the use of an in vitro cell culture system. Both heart-derived (Liu et al., 2020) and stem-cell differentiated (Ladha et al., 2021) cardiomyocytes in cell culture are known to exhibit immature phenotypes including Z-line disarray and TT disruption (Guo and Pu, 2020; Yang et al., 2014). Thus, whether proximity proteomic results from in vitro systems represent the Z-line/TT proteome in native cardiomyocytes remain questionable. Alternatively, transgenic or knock-in mice expressing BioID2 or APEX2 have been reported to enable in vivo proximity proteomics in the heart (Feng et al., 2020; Liu et al., 2020). However, in vivo proximity proteomics analysis is yet to be directly applied to the Z-line due to the absence of suitable mouse models.

To overcome the technical issues above, we recently harnessed adeno-associated virus (AAV) as a rapid and convenient method to perform cardiomyocyte-specific proximity labeling in mice. Using JMC markers Triadin and Junctin as baits, we successfully identified CMYA5 as a novel regulator of JMCs (Lu et al., 2022). Here, we established a new AAV-based ACTN2-BioID2 vector to perform in vivo proximity proteomics analysis of cardiomyocyte Z-lines in mice. We identified a novel Z-line-associated protein, palmdelphin (PALMD), that plays a critical role in modulating NEXN, Z-line-JMC tethering, and myocardial pathogenesis.

## Results

### In vivo proximity proteomics depicts a protein atlas of Z-lines in the heart

We constructed an AAV vector to express ACTN2-BioID2 fusion protein via a cardiomyocyte-specific *Tnnt2* promoter (Lian et al., 2012). We subcutaneously administered 5×10^10^ AAV to postnatal day 1 (P1) mice and intraperitoneally injected 24 mg/kg biotin twice a day for three days before cardiac ventricles were collected for further analysis at P14 (Fig. 1A). Western blotting showed that the AAV vector expressed ACTN2-BioID2 at a level that was much lower than endogenous ACTN2 in the heart (Fig. S1A), so that the ectopic expression was unlikely to perturb the heart or result in aberrant ACTN2 interactions. Immunofluorescence imaging of isolated cardiomyocytes demonstrated colocalization between the HA tag of ACTN2-BioID2 and cysteine and glycine⍰rich protein 3 (CSRP3), a classic Z-line marker (Rashid et al., 2015) (Fig. 1B). Fluorescent streptavidin also positively stained Z-lines (Fig. 1B), indicating specific proximity labeling of Z-line-associated proteins.

**Figure 1.**
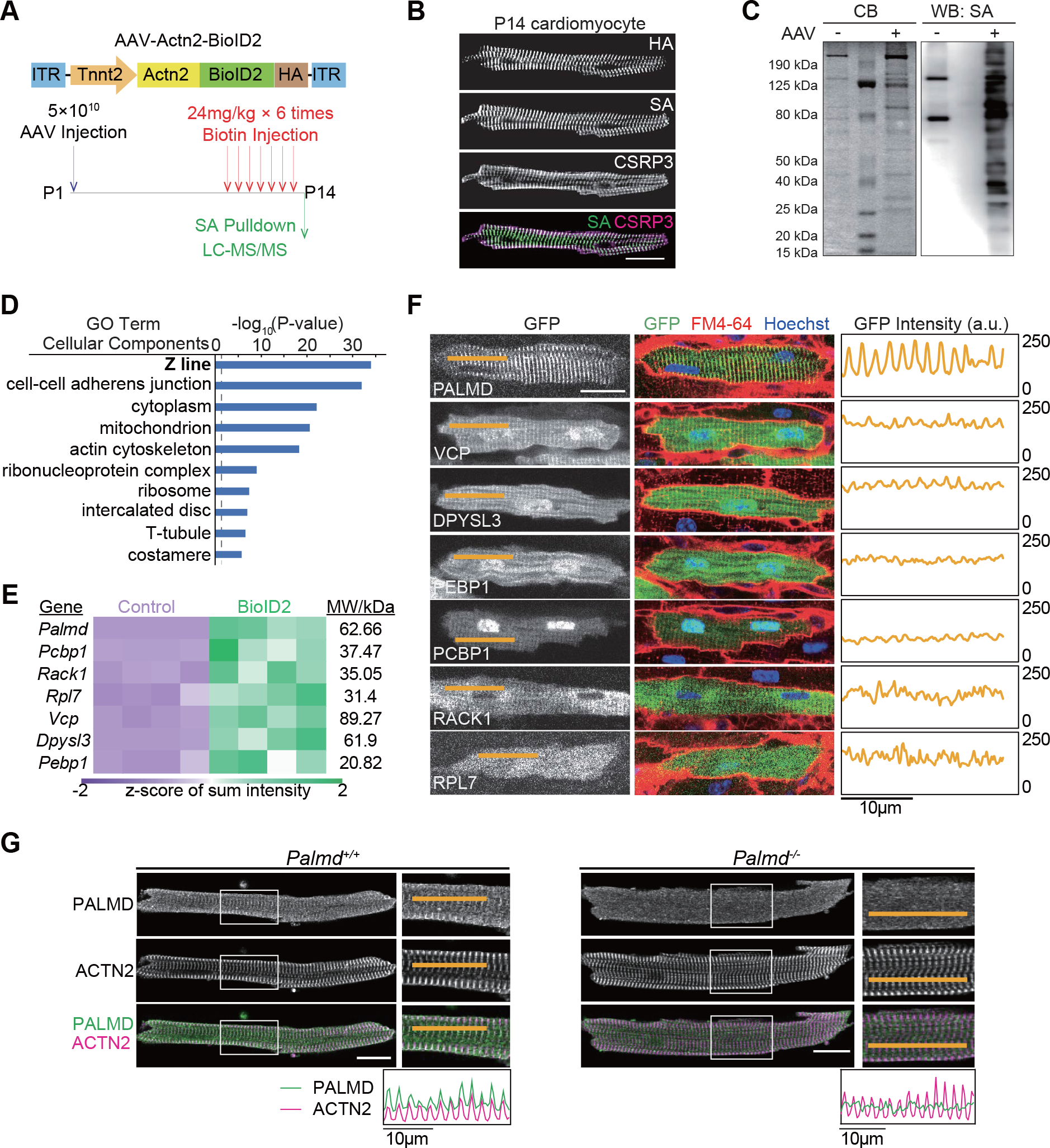
In vivo proximity proteomics uncovers PALMD as a novel Z-line-associated protein. (A) Diagram of the experimental workflow. Experimental and control hearts were collected from littermates that were injected with either AAV or saline. ITR, inverted terminal repeat. HA, hemagglutinin. SA, streptavidin. LC-MS/MS, liquid chromatography tandem mass spectrometry. (B) Immunofluorescence imaging of P14 isolated cardiomyocytes. (C) Coomassie blue and SA staining on a full-length SDS-PAGE blot. CB, coomassie blue. (D) The top 10 Gene Ontology Cellular Component terms that were significantly enriched in the ACTN2-BioID2 samples. P-value indicates a modified Fisher’s Exact P-value. (E) Heatmap of LC-MS/MS data for candidate proteins. MW, molecular weight. (F) In situ cardiac confocal imaging of GFP-tagged candidate proteins. The hearts were counterstained with plasma membrane/TT dye FM 4-64. Kymographs along the orange lines are shown to the right of the images. a.u., arbitrary unit. (G) Immunofluorescence imaging of isolated adult murine cardiomyocytes. Boxed areas were enlarged to the right of the original images. The intensity of PALMD and ACTN2 signals along the orange lines are depicted in the kymographs. Scale bar, 20 μm.

We next pulled down and purified the biotinylated proteins in cardiac extracts using streptavidin beads. Coomassie blue staining and streptavidin blots demonstrated that ACTN2-BioID2 indeed triggered the biotinylation of Z-line proteins that were absent in the control samples that did not receive AAV treatment (Fig. 1C). Endogenous ACTN2 and cardiac α-actin (ACTC1), which directly interact with ACTN2, were validated in the pulldown samples only when both AAV and biotin were given to the animals (Fig. S1A). Based on these validation data, the streptavidin beads were subjected to trypsin digestion followed by mass spectrometry-based protein analysis.

We compared the streptavidin pulldown proteomes between AAV-ACTN2-BioID2-treated samples and no AAV-treated control samples and identified 237 proteins that were significantly enriched in the ACTN2-BioID2 samples (Fig. S1B). Gene ontology (GO) analysis showed Z-line as the most enriched GO among Cellular Component (CC) gene sets (Fig. 1D). We manually curated each protein hit and found more than 20 proteins in our data that were previously reported to localize on Z-lines (Fig. S1C). Markers of TTs, costameres (containing adherent junction proteins), and intercalated discs were also detected as expected (Fig. 1D and S1C), since these subcellular structures are tethered to myofibrils at Z-lines.

Next we compared our data with published ACTN2-BioID data that were generated in hPSC-CMs (Ladha et al., 2021). We observed ∼41% overlap among identified proteins between the two data sets (Fig. S2A). These shared hits were enriched with classic Z-line, costamere, and intercalated disk markers (Fig. S1C and S2B), indicating that both approaches identify core Z-line-associated proteins. By contrast, the unique hits in the in vivo data were associated with mitochondria while the hPSC-CM data was more enriched with proteins in cytoplasm (Fig. S2B). These differences agreed with the observation that postnatal cardiomyocytes undergo massive biogenesis of mitochondria, which are associated with myofibrils in vivo, while hPSC-CMs exhibit immature phenotypes with a relatively larger fractional volume of cytoplasm.

### PALMD is a novel Z-line-associated protein in mouse and human

We next wondered if the ACTN2-BioID2 analysis could discover novel Z-line-associated proteins. We manually selected seven candidate proteins in the MS data that were not yet reported to be associated with Z-lines. In part, these proteins were selected because their coding sequences fit within the limited packaging capacity of AAV (Fig. 1E). We constructed AAV vectors that express each candidate fused to GFP. The AAV was injected into P1 pups. At P14, we performed in situ myocardial imaging of GFP, using FM 4-64 as a marker for TTs and Z-line (Chen et al., 2015; Guo et al., 2017). Among the seven candidates, palmdephin (PALMD) demonstrated the most prominent Z-line/TT localization (Fig. 1F). To a lesser extent, VCP, DPYSL3, PEBP1 and PCBP1 also exhibited Z-line-associated patterns.

PALMD belongs to the paralemmin protein family and was recently reported to be expressed in cardiomyocytes, endothelial cells and valvular interstitial cells in the heart (Sáinz-Jaspeado et al., 2021; Wang et al., 2022). Immunofluorescence analysis of PALMD in adult murine cardiomyocytes clearly demonstrated overlapping signals between PALMD and ACTN2 (Fig. 1G). A germline *Palmd* knockout (*Palmd*^*-/-*^) mouse was generated via zygotic CRISPR/Cas9 genome editing (Fig. S3A). Intercrosses bewteen *Palmd*^*+/-*^ mice yielded the expected Mendelian genotype distribution (Fig. S3B). *Palmd*^*-/-*^ mice exhibited no survival or growth defects for at least six months of age (Fig. S3C). Complete depletion of PALMD in the *Palmd*^*-/-*^ hearts was verified by Western blotting (Fig. S3D). In *Palmd*^*-/-*^ adult cardiomyocytes, Z-line PALMD immunostaining signals were ablated (Fig. 1G). These data validated endogenous PALMD as an authentic Z-line protein.

In the published ACTN2-BioID data in hPSC-CMs (Ladha et al., 2021), PALMD was not detected. To determine if this was a false negative result or reflects differences between human PSC-CMs versus mouse CMs. Thus, we derived cardiomyocytes from human embryonic stem cells following a protocol similar to the previous study (Ladha et al., 2021; Lian et al., 2012) and performed immunofluorescence analysis of PALMD on these cells. Interestingly, PALMD was observed to overlap with ACTN2 in ∼20% hPSC-CMs in sharp contrast to the ∼97% in adult murine cardiomyocytes (Fig. S4). These data indicate that PALMD is a Z-line protein in human cardiomyocytes, but the immature phenotypes of hPSC-CMs compromised the capacity of proximity proteomics to detect PALMD in hPSC-CMs.

### PALMD loss results in aggravated cardiac injury upon isoproterenol stimulation

PALMD modulates the actin cytoskeleton and contributes to aortic valve calcification (Li et al., 2020; Sáinz-Jaspeado et al., 2021; Thériault et al., 2018; Wang et al., 2022), but its role in cardiomyocytes has yet to be determined. Echocardiography of adult *Palmd*^*-/-*^ mice demonstrated no detectable defects in systolic function or left ventricular dimensions (Fig. S5A). Immunostaining or in situ imaging analysis detected no alterations in the organization of ACTN2 or TTs (Fig. S5B-C). Isolated cardiomyocytes also exhibited no changes of cardiomyocyte size or geometry (Fig. S5D). Therefore, PALMD is dispensable for basal cardiomyocyte ultrastructure and functions.

We next tested whether PALMD is required for normal cardiac responses to stress. Isoproterenol is an adrenergic receptor agonist that is frequently applied to animals to mimic stress-induced heart injury (Leenen et al., 2001). We administered isoproterenol daily to adult mice and performed echocardiogram weekly (Fig. 2A). Fractional shortening (FS) appeared more impaired in *Palmd*^*-/-*^ mice compared to wildtype controls (Fig. 2A). Diastolic left ventricular inner diameter (LVIDd) was unaltered, indicating lack of ventricular dilatation, while left ventricular posterior wall thickness (LVPWd) was *Palmd*^*-/-*^ mice due to attenuated isoproterenol-induced wall thickening (Fig. 2B).

**Figure 2.**
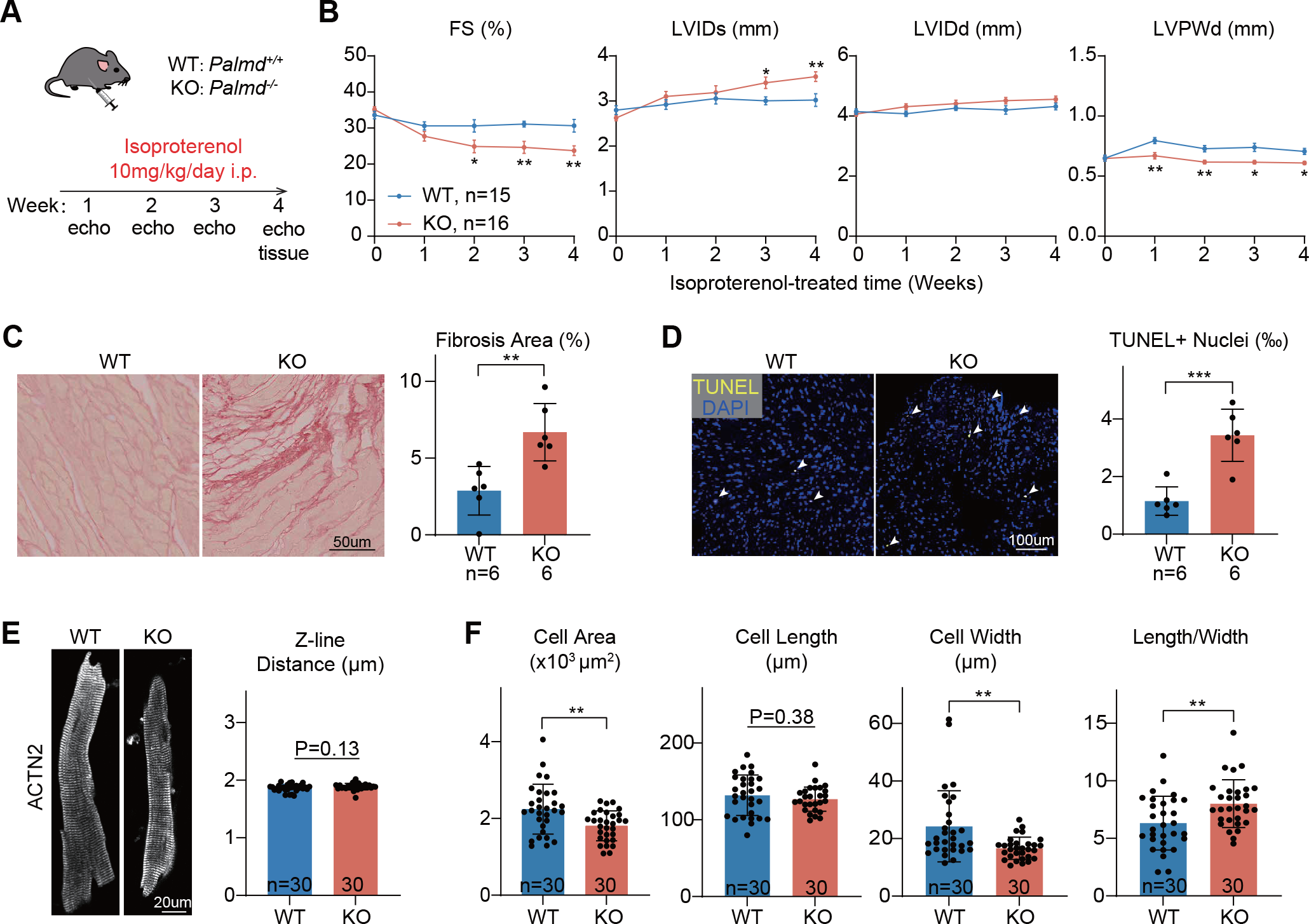
*Palmd* knockout aggravates isoproterenol-induced cardiac defects. (A) Diagram of the experimental workflow. i.p., intraperitoneal injection. (B) Echocardiography of cardiac function and dimensions. FS, fractional shortening. LVIDs, end systolic left ventricular internal diameter. LVIDd, end diastolic left ventricular internal diameter. LVPWd, end dyastolic left ventricular posterior wall thickness. (C) Picrosirius red staining of ventricle sections and quantification of fractional fibrosis area. (D) TUNEL staining on ventricle sections and quantification of TUNEL-positive nuclei. (E) Immunofluorescence images of ACTN2 in isolated cardiomyocytes and quantification of distance between adjacent Z-lines. Arrowheads point to TUNEL-positive nuclei. (F) Quantification of the size and morphology of isolated adult cardiomyocytes. Student’s t-test, *P<0.05, **P<0.01, ***P<0.001. In (B-D), n indicates animal numbers. In (E-F), n indicates cell numbers. In (B), mean ± SEM. In (C-F), mean ± SD.

Picrosirius red staining on cardiac sections indicated more severe cardiac fibrosis in *Palmd*^*-/-*^ hearts after isoproterenol treatment (Fig. 2C). Terminal deoxynucleotidyl transferase dUTP nick end labeling (TUNEL) demonstrated increased cell death in Palmd mutant hearts (Fig. 2D). In isolated cardiomyocytes, ACTN2 staining detected no changes of Z-line organization and distances (Fig. 2E). Projected cell area in mutant cardiomyocytes were less enlarged than control cells (Fig. 2F). Isoproterenol was known to induce concentric cardiac hypertrophy, which was characterized by cardiomyocyte widening and reduced length/width ratio (Nakamura and Sadoshima, 2018). In *Palmd*^*-/-*^ hearts,such cellular remodeling was compromised (Fig. 2F). Together, these data showed that PALMD depletion perturbed isoproterenol-induced cardiac hypertrophy and aggravate cardiac injury.

Because PALMD is expressed in both cardiomyocytes and non-cardiomyocytes in the heart (Sáinz-Jaspeado et al., 2021), we next tested if cardiomyocyte-specific PALMD depletion is sufficient to modify isoproterenol-triggered cardiac injury. We designed two sgRNAs targeting Palmd and constructed AAV vectors expressing sgRNA along with a Cre transgene controlled by a Tnnt2 promoter and a miR122 target sequence (miR122TS)-containing 3’ untranslated region (3’UTR)(Luzi Yang, 2023). This vector expresses Cre specifically in cardiomyocytes and enables cardiomyocyte specific *Palmd* somatic mutagenesis upon systemic AAV administration to Rosa^Cas9-tdTomato^ mice, which express Cas9 following Cre recombination (Guo et al., 2017; Lin et al., 2022) (Fig. S6A).

We first validated this Cas9/AAV9-based somatic mutagenesis (CASAAV) and observed robust cardiac *Palmd* gene mutagenesis and protein depletion by amplicon-sequencing and western blot, respectively (Fig. S6B-C). After isoproterenol treatment, CASAAV-treated mice exhibited more severe systolic dysfunction and compromised cardiac hypertrophy as compared to AAV-Cre treated controls (Fig. S6D). CASAAV-treated hearts also demonstrated increased interstitial fibrosis and cell death (Fig. S6E-F). Isolated cardiomyocytes exhibited size and morphology changes similar to the phenotypes in *Palmd*^*-/-*^ mice (Fig. S6G-H). No ACTN2 pattern or Z-line distance phenotypes were observed (Fig. S6G-H). Thus, loss of PALMD in cardiomyocytes sensitizes the heart to isoproterenol-induced injury and cardiac dysfunction.

### PALMD overexpression mitigates isoproterenol-induced cardiac injury

Next, we asked whether PALMD overexpression was sufficient to mitigate isoproterenol-induced cardiac injury. We harnessed the AAV-GFP-Palmd vector that was also used to validate PALMD Z-line localization (Fig. 1F) to overexpress PALMD using an AAV-GFP vector as control (Fig. 3A). 5×10^10^vg/g vectors were injected into P7 pups and daily isoproterenol injection was started at 4 weeks of age. Western blot analysis validated the overexpression of PALMD (Fig. 3B). Echocardiogram showed that Palmd overexpression protected mice from isoproterenol-induced systolic dysfunction (Fig. 3C). At the tissue level, the PALMD expression resulted in reduced cardiac fibrosis (Fig. 3D) and TUNEL-positive cell nuclei (Fig. 3E). While PALMD overexpression did not affect Z-line organization and spacing (Fig. 3F), it triggered more pronounced cardiomyocyte enlargement, mainly due to cellular widening and reduced length/width ratio (Fig. 3G). Together, these data implied that the AAV-based PALMD overexpression protects the myocardium during isoproterenol-induced cardiac injury.

**Figure 3.**
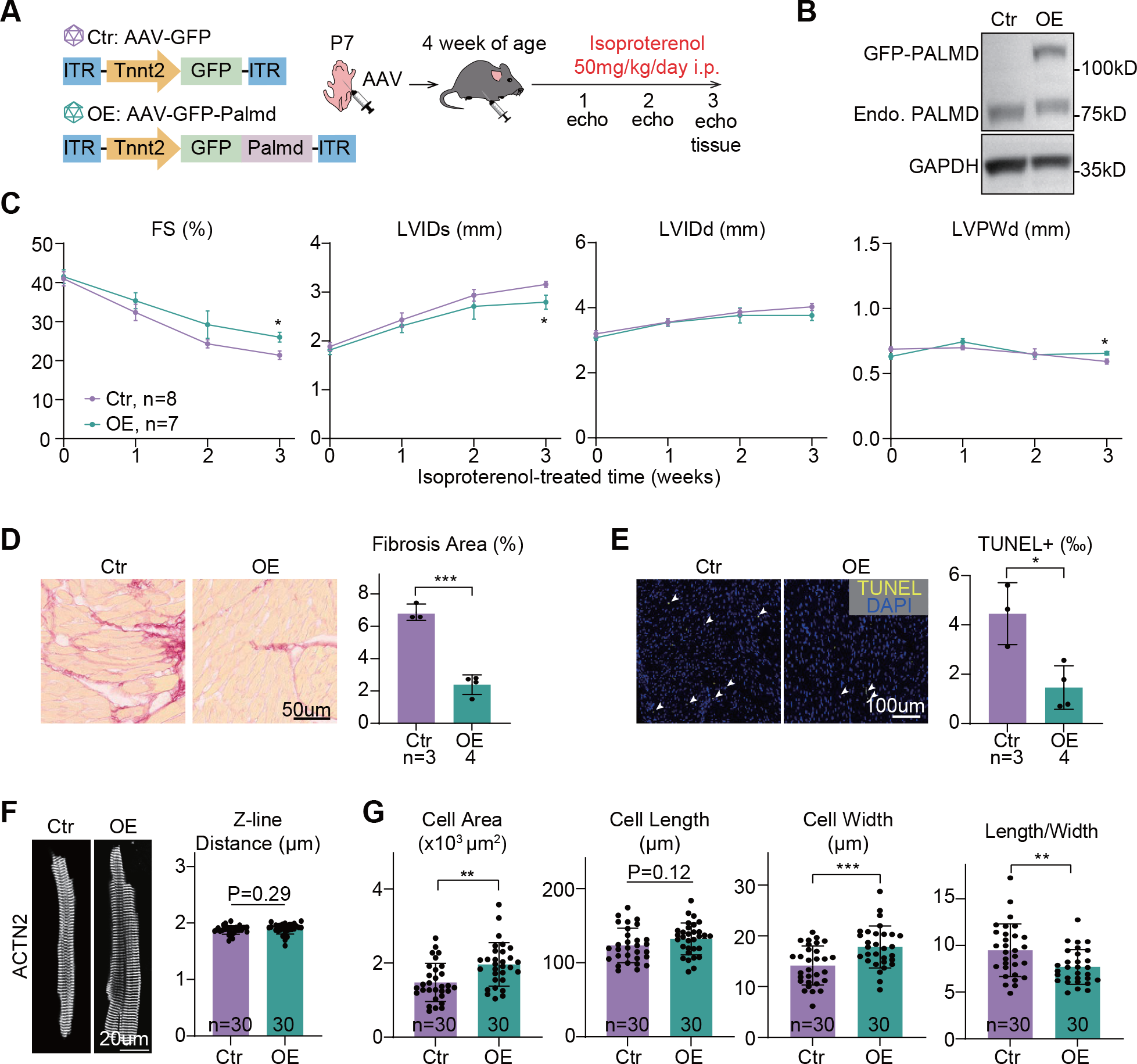
PALMD overexpression alleviates isoproterenol-induced defects. (A) Diagram of the experimental workflow. Ctr, control. OE, overexpression. (B) Western blot analysis of PALMD overexpression. (C) Echocardiography of cardiac function and structure. (D) Picrosirius red staining and quantification. (E) TUNEL staining and quantification. Arrowheads point to TUNEL-positive nuclei. (F) Immunofluorescence images of ACTN2 and quantification adjacent Z-line distance. (G) Quantification of the size and morphology of isolated adult cardiomyocytes. Student’s t-test, *P<0.05, **P<0.01, ***P<0.001. In (C-E), n indicates animal numbers. In (F-G), n indicates cell numbers. In (C), mean ± SEM. In (D-G), mean ± SD.

### PALMD stabilizes NEXN on Z-line and regulates JMCs

We next explored the potential mechanism by which PALMD regulated cardiac functions. Since PALMD depletion resulted in systolic dysfunction upon isoproterenol treatment, we first tested an array of known Z-line-associated proteins that were required to maintain systolic function. We examined the Z-line localization of CSRP3, LIM domain binding 3 (LDB3), palladin (PALLD), myopalladin (MYPN), filamin C (FLNC), and nexilin (NEXN) in isoproterenol-treated *Palmd-/-* cardiomyocytes, and we found the Z-line pattern of nexilin (NEXN) was largely diminished upon PALMD depletion (Fig. 4A and S7A). The NEXN content was analyzed by AutoTT, a software that objectively quantifies T-tubule and other sarcomere system contents by normalizing both transversely oriented and longitudinally oriented TT-like patterns to cell morphology(REF). Western blot analysis demonstrated significantly reduced NEXN protein levels (Fig. 4B) while the *Nexn* mRNA level was not influenced (Fig. S7B), suggesting a post-transcriptional mechanism. In a co-immunoprecipitation assay in HEK293T cells, HA-PALMD and FLAG-NEXN could mutually pull down each other (Fig. 4C), indicating interaction between these two proteins. In NEXN-expressing cells treated with 20 μg/ml cycloheximide, a protein translation inhibitor, we observed more profound decrease of NEXN in the absence of PALMD (Fig. 4D), further confirming a critical role of PALMD in stabilizing NEXN protein.

**Figure 4.**
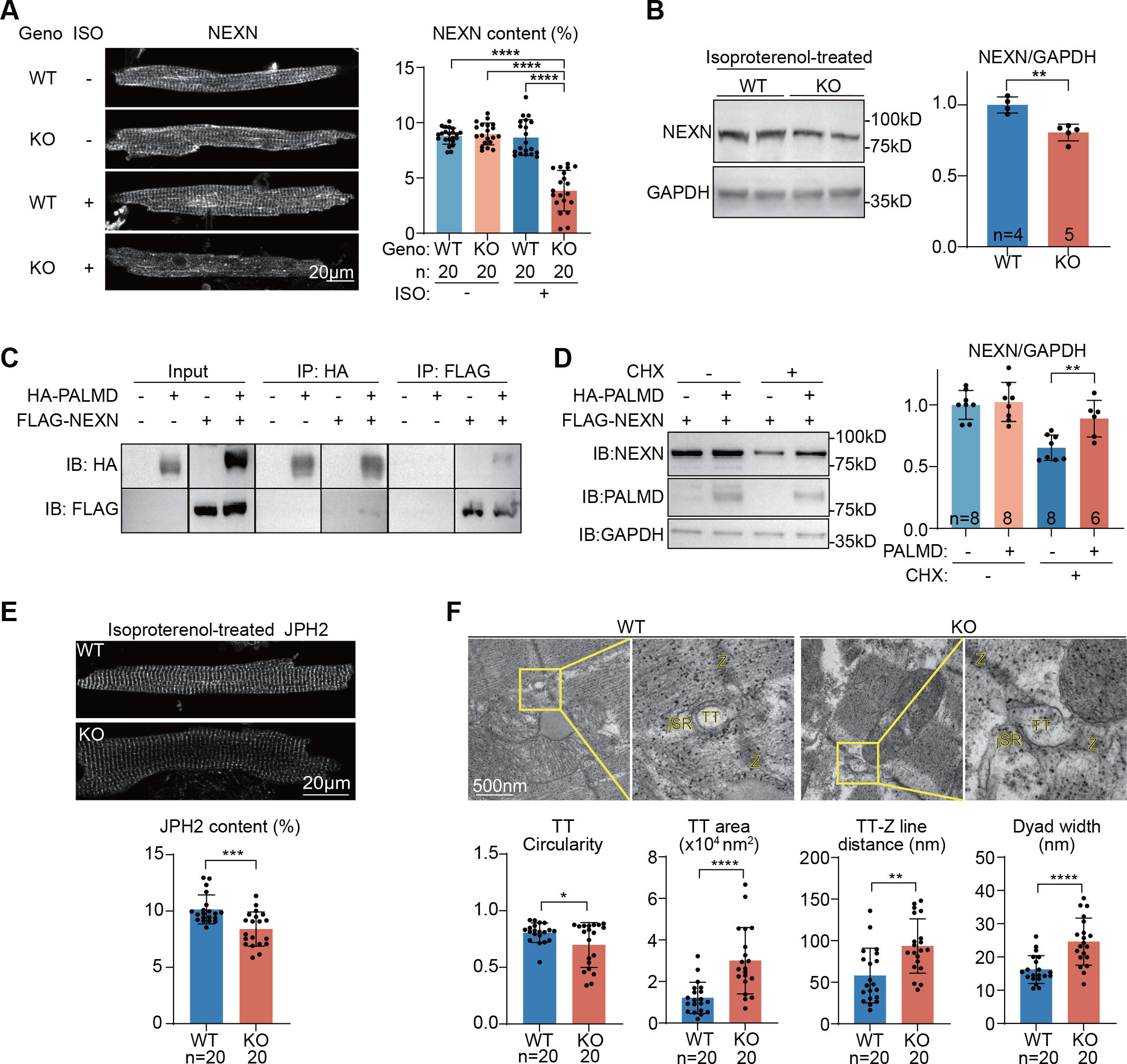
PALMD modulates JMCs by stabilizing NEXN. (A) Immunofluorescence imaging and AutoTT quantification of NEXN patterns on Z-line. ISO, isoproterenol. (B) Western blots and quantification of NEXN protein in isoproterenol-treated hearts. (C) Co-immunoprecipitation analysis between HA-PALMD and FLAG-NEXN in HEK293T cells. IP, immunoprecipitation. IB, immunoblotting. (D) Western blot analysis and quantification of NEXN in cells treated with 20 μg/ml cycloheximide for 12h. (E) Immunofluorescence imaging and AutoTT quantification of JPH2 patterns on Z-line. (F) Transmission electron microscopy analysis of junctional membrane complexes and quantification. TT, transverse tubule. jSR, junctional sarcoplasmic reticulum. Z, z-line. Student’s t-test, *P<0.05, **P<0.01, ***P<0.001, ****P<0.0001. In (A, E), n indicates cell numbers. In (B), n indicates animal number. In (D), n indicates repeated experiments. In (F), n indicates JMC number. Mean ± SD.

NEXN was recently shown to interact with components in JMCs and regulated TT and JMC organization (Liu et al., 2019). Thus, we next examined the role of PALMD in stabilization of JMCs and TTs. Immunofluorescence of junctophilin-2 (JPH2), a structural protein that tethers TT and SR membranes at JMCs (Landstrom et al., 2014), uncovered significantly reduced Z-line pattern of JPH2 in PALMD-depleted cardiomyocytes, as measured by the AutoTT software (Guo and Song, 2014) (Fig. 4E). A similar phenotype was also detected for another TT marker caveolin-3 (CAV3) (Fig. S7C). No changes of JPH2 or CAV3 protein levels were detected by western blot (Fig. S7D). By transmission electron microscopy, PALMD depletion also resulted in reduced TT circularity, increased TT luminal area, increased TT distances to Z-line and SR membranes (Fig. 4F). Together, these data showed that PALMD plays a crucial role in maintaining JMCs and their association with Z-line in the isoproterenol-stressed conditions through stabilizing NEXN.

## Discussion

Z-lines are core ultrastructural organizers of cardiomyocytes that modulate many facets of cardiac pathogenesis. Comprehensive characterization of Z-line proteins is pivotal in the efforts to dissect molecular mechanisms and identify potential therapeutic targets for heart diseases. ACTN2-based proximity proteomics was recently reported in hPSC-CMs (Ladha et al., 2021). However, because hPSC-CMs in cell culture are known to be structurally immature, exhibiting disrupted Z-line and TTs (Guo and Pu, 2020; Yang et al., 2014), there are potential advantages to performing these studies in de facto cardiomyocytes that have normal Z-line structure and organization.

In this study, we established an AAV-delivered, cardiomyocyte-specific, proximity-labeling approach to characterize the Z-line-associated proteome in vivo. We discovered an array of novel Z-line-associated proteins and further investigated a new role of PALMD in cardiac pathogenesis. The Z-line-associated localization of PALMD was confirmed in both murine cardiomyocytes and hPSC-CMs, but was undetected in the previous proximity proteomics data in hPSC-CMs, suggesting that the immature phenotypes of hPSC-CMs limited the set of Z-line-associated proteins detected by proximity-labeling.

PALMD was reported to be expressed in multiple cell types in the heart. However, the function of PALMD in cardiomyocytes was unclear. In this study, we discovered PALMD to be dispensable for cardiac homeostasis in the unstressed heart. However, isoproterenol stress, which models cardiac injury caused by chronic sympathetic activation in heart disease, PALMD depletion resulted in aggravated cardiac injury and dysfunction. Overexpression of PALMD partially mitigated isoproterenol-induced cardiac injury, nominating PALMD as a novel therapeutic target for cardiac protection. Strikingly, we found that PALMD played a role in stabilizing NEXN, a critical Z-line-associated protein that was reported to regulate both Z-line stability and TT formation. Thus, PALMD participates in the crosstalk between Z-lines and TTs/JMCs mediated by NEXN. Mutations of many Z-line components, including NEXN, are known to cause inherited cardiomyopathy. Therefore, further investigation of the mechanisms by which PALMD regulates NEXN under the stressed conditions would likely uncover novel molecular mechanisms behind cardiomyopathy pathogenesis from the standpoint of Z-line-JMC crosstalk.

The disassembly of Z-lines is a hallmark of cardiomyocyte dedifferentiation and has been observed in cardiomyocyte regeneration as well as many forms of heart diseases (Zhu et al., 2021). The AAV-ACTN2-BioID2 approach may offer a new inroad toward dissection of cardiomyocyte dedifferentiation. Future investigation of the PALMD-NEXN axis in other forms of heart diseases and cardiac regeneration would be intriguing. In addition, further mining of our proximity proteomics data and research on other new Z-line proteins identified in this study will yield deeper understanding of Z-line-based regulation of cardiac pathogenesis.

## Author Contributions

W.T.P. and Y.G. provided the overall supervision; Y.G. conceived the research; C.G. performed major experiments and analysis of Palmd subcellular localization and function in the heart; B.D.J. and R.A conducted BioID experiments; J.L. performed echocardiogram analysis; L.Y. upgraded the CASAAV vectors and managed the Rosa^Cas9-Tom^ mice; F.L. provided critical guidance on BioID; Q.M and N.M. managed mouse husbandry and experiments; Y.C. assisted in data collection and analysis; C.L. provided critical guidance on NEXN analysis; Z.W., X.L., and F.L. conducted hPSC-CM analysis; M.Z., H.X. and E.D. offered help in isoproterenol treatment experiments; Y.G. and C.G. wrote the manuscript; W.T.P edited the manuscript.

## Acknowledgments

We appreciate Taplin Mass Spectrometry Facility at Harvard Medical School for mass spectrometry analysis, Mouse Gene Manipulation Core at Boston Children’s Hospital for the generation of *Palmd*^*-/-*^ mice, and PackGene Biotech for AAV packaging.

## Funding information

Y.G. was funded by the National Key R&D Program of China (2022YFA1104800), the National Natural Science Foundation of China (82222006, 32100660 and 82170367), Beijing Nova Program (Z211100002121003 and 20220484205) and Beijing Natural Science Foundation (7232094). W.T.P. was supported by National Institute of Health (R01HL146634 and R01HL163937). E.D.D. was supported by the Natural Science Foundation of China (82070235 and 92168113), the CAMS Innovation Fund for Medical Sciences (2021-I2M-5-003) and Haihe Laboratory of Cell Ecosystem Innovation Fund (HH22KYZX0047). C.L. was supported by the National Natural Science Foundation of China (82170231) and Guangdong Basic and Applied Basic Research Foundation (2023B1515020103).

## Competing Interest Statement

The authors declare no competing interests. A patent was filed to protect the AAV-ACTN2-BioID2 vector and its applications.

## Resource availability

### Lead contact

Further information and requests for resources and reagents should be directed to and will be fulfilled by the lead contact, Dr. Yuxuan Guo.

## Materials and data availability

Plasmids generated in this study will be available at Addgene. This study did not generate new unique reagents.

Other materials generated in this study are available from the Lead Contact upon reasonable request.

## Data and code availability

Raw mass-spectrometry data were attached in Table S1.

No unique code was generated for the analysis.

Any additional information required to reanalyze the data reported in this paper is available from the Lead Contact upon reasonable request.

## Experimental model and subject details

### Animal

Animal strains and procedures were approved by the Institutional Animal Care and Use Committee of Boston Children’s Hospital (19-07-3969R) and Peking University (LA2021332). The *Rosa*^*Cas9-Tom*^ mice were purchased from GemPharmatech (Strain No. T002249) and validated as previously described (Lin et al., 2022). The *Palmd*^*-/-*^ mice were generated at Mouse Gene Manipulation Core at Boston Children’s Hospital via CRISPR/Cas9-based zygotic mutagenesis. See Table S2 for sgRNA and genotyping information. Animals are housed at the Peking University Health Science Center Department of Laboratory Animal Science and are regularly monitored with respect to general health, cage changes, and overcrowding. Littermates of the same sex were randomly assigned to experimental groups.

### hPSC-CM differentiation

The hESC line MEL-1 (NIHhESC-11-0139) were maintained in PGM1 hPSC culture medium (Cellapy #CA1007500, China) on plates coated with PSCeasy coating solution (Cellapy #CA3003100, China). The cells were maintained at 37°C in humidified air with 5% CO_2_. The cells were passaged at a 1:6 ratio with 0.5 mM EDTA/PBS when 80%-100% confluency was achieved.

MEL-1 cells were differentiated into cardiomyocytes by the CardioEasy human cardiomyocyte differentiation kit (Cellapy #CA2004500, China) using an established protocol (Lian et al., 2012). In brief, when hPSCs reached 80%-95% confluency at day 0, the medium was replaced with CardioEasy differentiation medium 1. On day 2, the medium was replaced with CardioEasy differentiation medium 2. From day 4 onward, the medium was replaced with CardioEasy complete medium every other day. On day 30, cells were purified using CardioEasy cardiomyocyte purification kit (Cellapy #CA2005100, China) using an established protocol(Tohyama et al., 2016). After purification, cells were maintained in CardioEasy complete medium. The differentiated cells were digested with CardioEasy human cardiomyocyte digestion kit (Cellapy #CA2004500, China) and seeded onto glass-bottom dishes before immunofluorescence.

## Method details

### Plasmid

The BioID2-HA coding sequence was acquired from Addgene (#80899)(Kim et al., 2016) and subcloned into our previously published AAV-cTNT-Actn2-GFP vector (Addgene #165034) (Guo et al., 2021) to generate the AAV-cTNT-Actn2-BioID2-HA vector. Coding sequences for *Rpl7* (MMM1012-202805808) were from Dharmacon, USA. *Vcp* (MR210760), *Dpysl3* (MR220351), *Pebp1* (MR201759) and *Rack1*(MR204575) cDNA clones were purchased from Origene, USA. *Palmd* and *Pcbp1* were cloned from heart cDNA. They were subcloned into the AAV-cTNT-Actn2-GFP vector to generate GFP-tagged gene overexpression vectors.

We incorporated a miR122TS sequence into our original CASAAV vector (Addgene # #87682) (Guo et al., 2017) to generate the AAV-U6-sgRNA1-U6-sgRNA2-cTNT-Cre-miR122TS vector. This new design could reduce leaky gene editing in the liver and greatly enhanced the cardiac specificity of in vivo genome editing (Luzi Yang, 2023). Next, we designed sgRNAs targeting *Palmd* using CRISPick (portals.broadinstitute.org/gppx/ crispick/public) and constructed CASAAV plasmids as previously described. See Table S2 for sgRNA information.

For co-immunoprecipitation experiments, the *Palmd* coding sequence was subcloned into our previously published pHA-ACTC1-R64D plasmid to generate the pHA-Palmd plasmid. The Nexn coding sequence was PCR amplified from the cDNA of murine hearts and subcloned into pFLAG-MRTFA (Addgene #11978) to create the pFLAG-Nexn plasmid.

### AAV production

AAV9 was prepared as previously described (Guo et al., 2017). AAV titer was quantified by real-time quantitative PCR. See Table S2 for primer information. Alternatively, AAV9 was packaged at PackGene Biotech. AAV was injected into P1 pups subcutaneously, or administered into P7 pups via Intraperitoneal Injection. The pups were anesthetized in an isoflurane chamber before injection.

### BioID and Mass Spectrometry

Animals were subcutaneously injected with AAV at P1. 24 mg/kg bodyweight biotin was intraperitoneally injected into mice twice a day for 3 consecutive days before sample collection. 2-3 hearts were pooled in a sample. Heart tissues were homogenized with RIPA buffer containing 1% SDS with protease and phosphatase inhibitors and centrifuged at >13000g for 10 mins to collect supernatant for BioID. Biotinylated proteins were enriched using magnetic streptavidin beads (Dynabeads M-280, Invitrogen, USA). The beads were washed with a series of SDS buffer, high salt buffer, LiCl buffer and stored in PBS at -80°C. BioID samples were validated by Simply Blue Stain (Life Technologies/Invitrogen LC6060) before mass spectrometry was performed.

Liquid chromatography with tandem mass spectrometry was performed at the Taplin Biological Mass Spectrometry Facility, Harvard Medical School, as previously described(Lu et al., 2022). In brief, 200□ng/μL sequencing-grade trypsin (Promega, Madison, WI) was incubated with the samples at 37□°C overnight. The beads were then removed using a magnet, and the supernatant was dried in a speed-vac. The samples were re-suspended in HPLC solvent A (2.5% acetonitrile, 0.1% formic acid). A nano-scale reverse-phase HPLC capillary column was created by packing 2.6□μm C18 spherical silica beads into a fused silica capillary (100□μm inner diameter□×□∼30□cm length) with a flame-drawn tip. After equilibrating the column, each sample was loaded via a Famos autosampler (LC Packings, San Francisco CA). A gradient was formed, and peptides were eluted with increasing concentrations of solvent B (97.5% acetonitrile, 0.1% formic acid). As peptides eluted, they were subjected to electrospray ionization and then entered an LTQ Orbitrap Velos Elite ion-trap mass spectrometer (Thermo Fisher Scientific, Waltham, MA). Peptides were detected, isolated, and fragmented to produce a tandem mass spectrum of specific fragment ions for each peptide. Peptide sequences (and hence protein identity) were determined by matching protein databases with the acquired fragmentation pattern by the software program, Sequest (Thermo Fisher Scientific, Waltham, MA). All databases include a reversed version of all the sequences and the data was filtered to between a one and two percent peptide false discovery rate.

### Echocardiography

Echocardiography was measured on mice that were anesthetized initially by 3% isoflurane and maintained asleep at 1∼1.5% isoflurane. Echocardiography was performed with a VINNO6n machine (VINNO Corporation, Suzhou, China). The standard long-axis cardiac videos were acquired using a 23MHz transducer. FS, LVPW and LVID values were measured by averaging results from five consecutive heart beats. The researcher who performed echocardiography was blinded to mouse genotypes.

### Amplicon sequencing

Genomic DNA was extracted from tissues using TIANamp Genomic DNA Kit (DP304, Tiangen, China). The sgRNA-targeted loci of Palmd gene were amplified using Taq PCR MasterMix (KT211, Tiangen, China) and purified by TIANgel Purification Kit (DP219, TIANGEN). See Table S2 for primer sequences. Sequencing was performed on an Illumina NovaSeq 6000 platform at Novogene, China. The sequencing results were processed by CRISPResso2(Clement et al., 2019).

### RNA extraction and RT-qPCR

Total RNA was extracted from heart apex using the TransZol Up Plus RNA Kit (ER501-01, TransGen, Beijing, China). TransScript II One-Step gDNA Removal and cDNA Synthesis SuperMix (AH311-03, TransGen, Beijing, China) kits were used for genomic DNA removal and reverse transcription. Real-time PCR analysis was performed using Perfect Start Green qPCR Super Mix (+DyeII) (AQ602-24, TransGen, Beijing, China) and the AriaMx Real-Time PCR System (Agilent Technologies). See Table S2 for primer information.

### In situ cardiac confocal imaging

Hearts were perfused with FM 4-64 (Invitrogen, 13320) on a Langendorff apparatus for 10 min at RT. The hearts were then positioned on a glass-bottom dish and immediately imaged using a Leica TCS SP8 MP FLIM inverted confocal microscope.

### Cardiomyocyte isolation

Cardiomyocytes were isolated by retrograde perfusion. In brief, heparin-treated mice were anesthetized with 3% isoflurane. Hearts were extracted and cannulated onto a Langendorff perfusion apparatus. 37°C perfusion buffer was first pumped into the heart to flush out blood and equilibrate the heart. Collagenase II (Worthington, LS004177) was next perfused into the heart for 8 min at 37 °C to dissociate cardiomyocytes. The apex was cut from the digested heart, gently dissociated into single cardiomyocytes in 10% FBS/perfusion buffer and filtered through a 100μm cell strainer to remove undigested tissues.

### Fibrosis analysis

Paraffin sectioning was performed at Servicebio, China. Heart tissues were fixed in 4% paraformaldehyde (PFA) at 4°C overnight and then in 20% sucrose solution overnight. Next, the samples were dehydrated through a serial gradient of ethanol and N-butanol. The samples were waxed in liquid paraffin for 2 h and embedded to make paraffin blocks using a tissue embedder (EG1150, Leica, Germany). Four-micron sections were cut by paraffin slicer (RM2245, Leica, Germany) and adhered to the microscope slides after floating on a water bath (HI1210, Leica, Germany).

For picrosirius red staining, the paraffin sections were dewaxed and dehydrated and then incubated with 0.2% picrosirius red solution dissolved in saturated aqueous picric acid (1.2% picric acid in water) (G1018, Servicebio). Next the sections were rinsed in water, dehydrated in increasing concentrations of ethanol, cleared in xylene and mounted for imaging using a Grundium Ocus^®^40 slides scanner.

### TUNEL analysis

Paraffin sectioning was performed at Servicebio, China. Heart tissues were fixed in 4% paraformaldehyde (PFA) at 4°C overnight and then in 20% sucrose solution overnight. Next, the samples were dehydrated through a serial gradient of ethanol and N-butanol. The samples were waxed in liquid paraffin for 2 h and embedded to make paraffin blocks using a tissue embedder (EG1150, Leica, Germany). Four-micron sections were cut by paraffin slicer (RM2245, Leica, Germany) and adhered to the microscope slides after floating on a water bath (HI1210, Leica, Germany).

For TUNEL assay, the paraffin sections were deparaffinized and rehydrated and then incubated with protease K working solution (G1205, Servicebio) at 37 °C to perform Antigen retrieval. Next the sections were permeabilized with permeabilize working solution (G1204, Servicebio) and equilibrium at room temperature. Tunel reaction was performed using tunel assay kit (G1501, Servicebio; G1502, Servicebio). The sections were then counterstained with DAPI (G1012, Servicebio) and mounted for imaging using a Keyence BZ-X810 all-in-one fluorescence microscope.

### Immunofluorescence analysis

Cardiomyocytes were fixed with 4% paraformaldehyde, permeabilized by 0.1% Triton-100/PBS and blocked in 4% BSA/PBS. Then the cells were incubated with primary antibodies overnight at 4 °C and then incubated with secondary antibodies and/or dyes at RT for 2 h. The cells were mounted with ProLong Diamond antifade mountant (Invitrogen, 36961) before imaging. See Table S3 for antibody and dye information.

Confocal images were taken using a Leica TCS SP8 MP FLIM laser-scanning confocal microscope with a 40X/1.5 objective. AutoTT (Guo and Song, 2014) was used to quantify T-tubule, JPH2 and NEXN. Cell size and shape were manually measured using ImageJ.

### Western blot analysis

Heart tissues were homogenized in RIPA buffer (25 mM Tris pH7.4, 150 mM NaCl, 1% Triton X-100, 0.5% Na Deoxycholate, 0.1% SDS) supplemented with protease inhibitor cocktail. Heart lysates were denatured in 2X SDS sample buffer at 70 °C for 10 min, separated on a 10% Bis-Tris gradient gel (Sangon Biotech, C691101-0001), transferred to a PDVF membrane, and blocked by 4% milk/TBST. Primary antibodies were incubated with the membrane overnight at 4°C. HRP-conjugated secondary antibodies were probed for 1∼2h at RT. Chemiluminescence was detected using an Invitrogen iBright™ CL1500. See Table S3 for antibody and dye information.

### Transmission electron microscopy

TEM experiments were performed by the Electron Microscopy Analysis Laboratory in Medical and Health Analysis Center, Peking university Health Science Center. Briefly, heart samples were collected, cut into small pieces (1–2 mm cubes), and fixed in EM Grade 2.5% Glutaraldehyde (P1126, Solarbio) overnight at 4 °C. Ultrathin sections were cut using an ultramicrotome (EM UC7, Leica, Germany) and stained with uranyl acetate/lead citrate. TEM images were obtained using a transmission electron microscope (JEM-1400PLUS, Leica, Germany). All the quantifications were done using ImageJ software.

## Quantification and statistical analysis

Statistics were performed using Prism (GraphPad) software. Imaging experiments were analyzed using ImageJ software. Data are presented as mean ± SD unless otherwise noted. Statistical analysis of all data was determined using the Student’s t test for two group comparisons and one-way ANOVA for comparison of multiple groups. Statistical significance was defined by *P<0.05, **P<0.01, ***P<0.001, ****P<0.0001.

## Graphical abstract

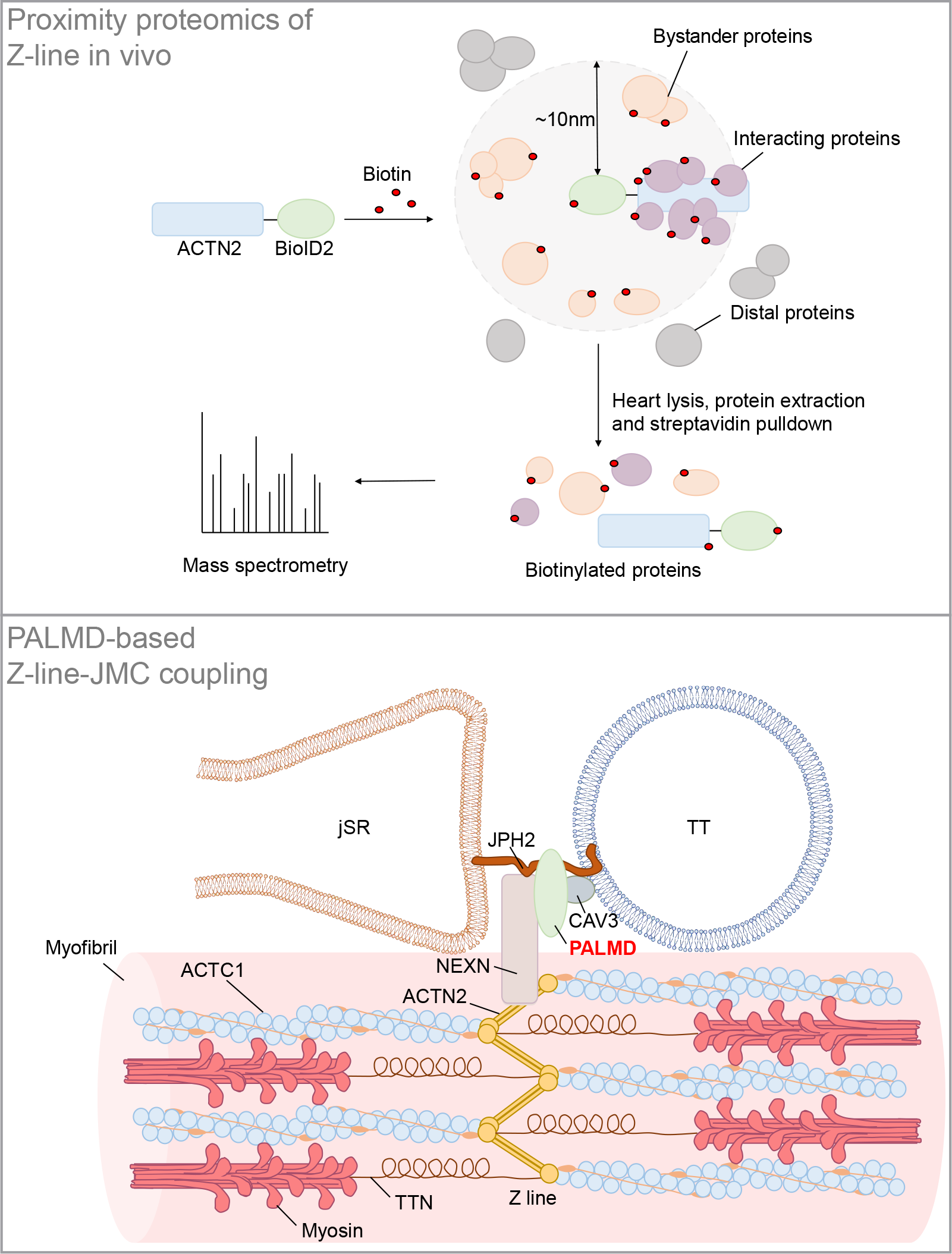

## Notes

### Competing Interest Statement

The authors have declared no competing interest.

