## Supplemental Data 1 for "In vivo proximity proteomics uncovers palmdelphin (PALMD) as a Z-line-associated mitigator of isoproterenol-induced cardiac injury"

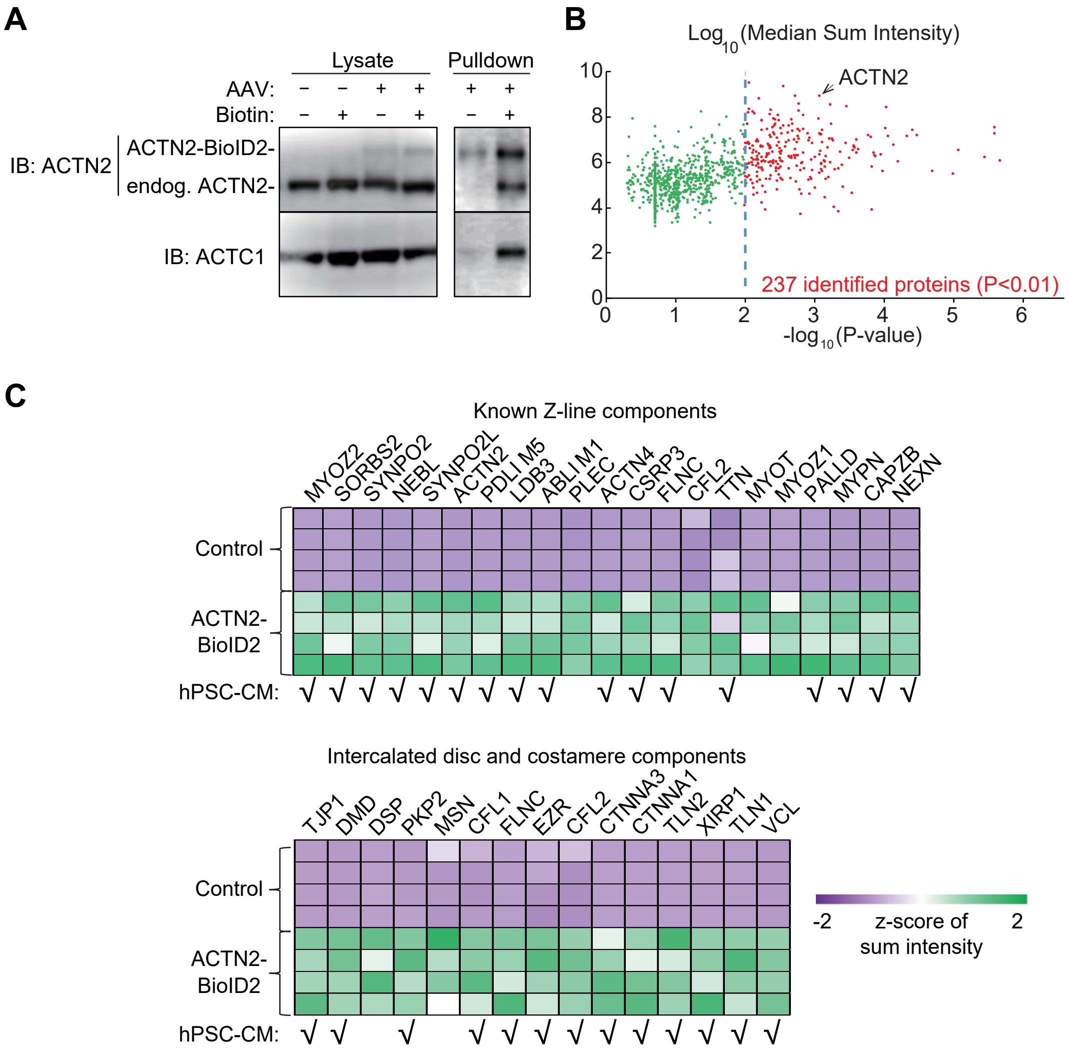


**Figure S1. Verification of ACTN2-BioID2-mediated in vivo proximity proteomics.** (A) Western blotting of cardiac lysates (left) and SA-pulldown proteins (right). Endog., endogenous. (B) A plot of MS sum intensity against P values for all identified proteins by ACTN2-BioID2 proteomics in vivo. P values were generated by student's t-test corrected by the Bonferroni's method. The dashed line indicates the cutoff for positive hits. (C) Heatmap of representative proteins in the MS data. Shared hits found in the previously published hiPSC-CM BioID experiment were checked below the heatmap.

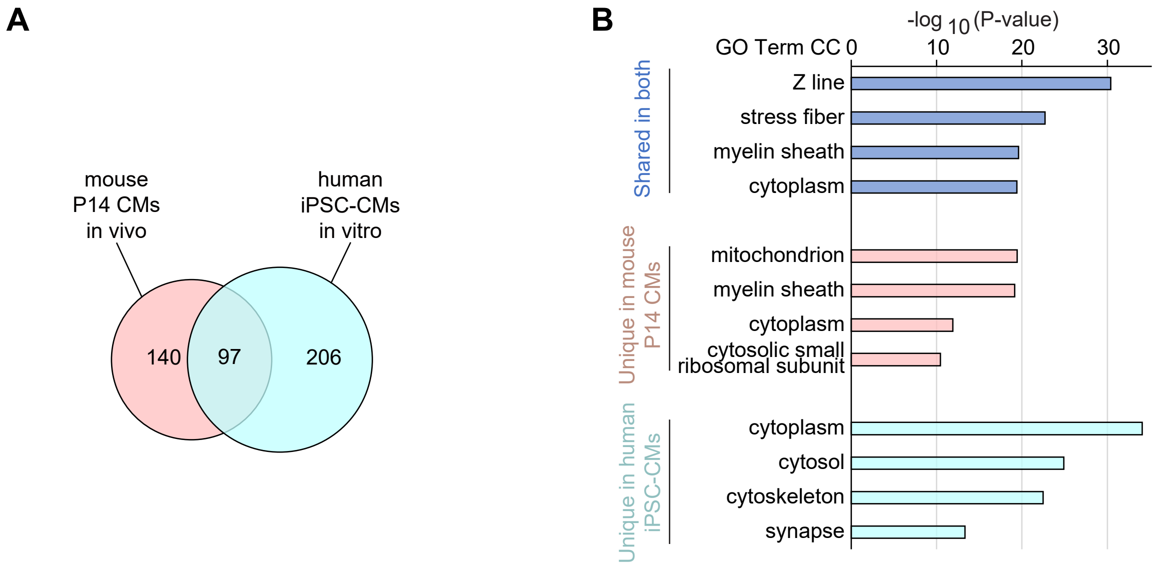


**Figure S2. Comparative analysis of ACTN2-BioID data in mice versus in hPSC-CMs.** (A)Venn diagram between ACTN2-BioID data in this study versus that in the previous hiPSC-CM study. (B) Top GO_CC terms enriched in the shared and unique ACTN2-BioID protein hits.

**A**

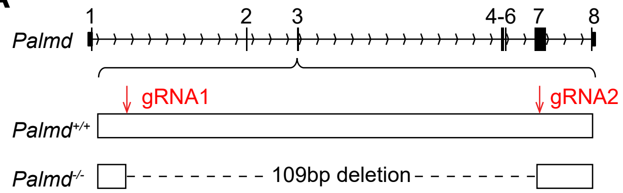


**B**

Genotype distribution at birth

| | +/+ (%) | +/- (%) | -/- (%) |
|---|---|---|---|
| Observed | 34 (29.0) | 58 (49.6) | 25 (21.4) |
| Expected | 29 (25.0) | 59 (50.0) | 29 (25.0) |

Pearson’s chi-squared test, $X^2$=0.702, P=0.704.

**C**

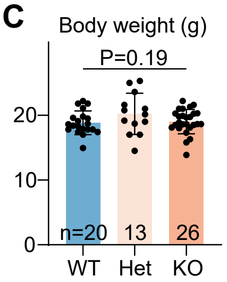


**D**

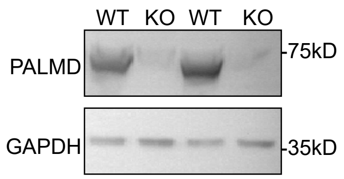


**Figure S3. Generation and basic characterization of Palmd$^{-/-}$ mice.** (A) A diagram showing sgRNA design inducing the actual mutation in the Palmd$^{-/-}$ mice. (B) Statistical analysis of offspring genotype distribution after Palmd$^{+/-}$ intercrosses. (C) Body weight analysis of 8~12-week-old animals. WT, Palmd$^{+/+}$. Het, Palmd$^{+/-}$. KO, Palmd$^{-/-}$.

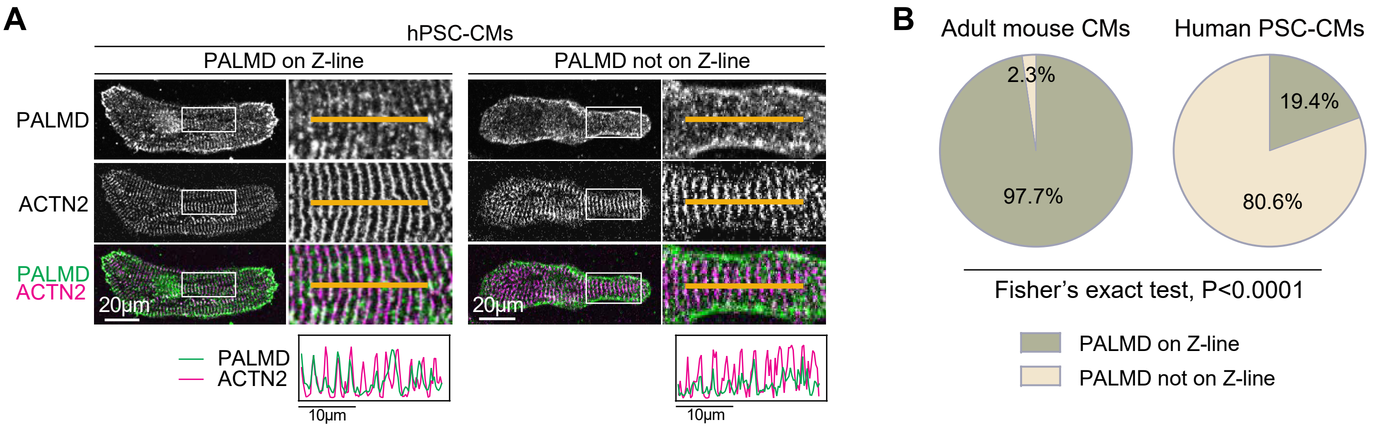


**Figure S4. PALMD subcellular localization in hPSC-CMs.** (A) Immunofluorescence images of PALMD in hPSC-CMs. (B) Quantification of cell fractions with PALMD on Z-lines in adult murine cardiomyocytes (n=43 cells) versus hPSC-CMs (n=98 cells).

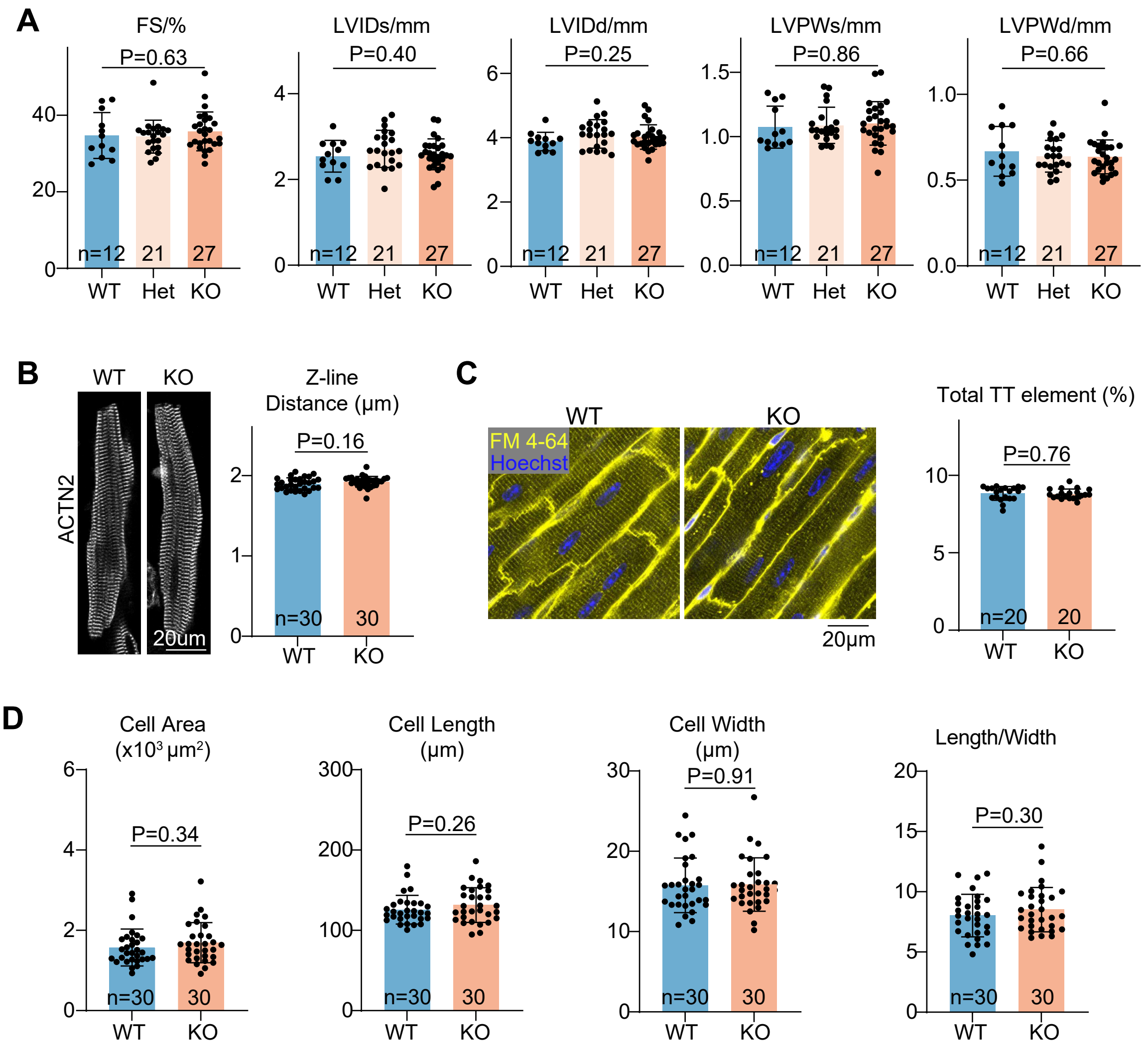


**Figure S5. Cardiac phenotypes in basal conditions.** (A) Echocardiogram analysis. (B) ACTN2 staining on isolated cardiomyocytes and Z-line distance quantification. (C) In situ myocardial imaging of FM 4-64-stained hearts and T-tubule quantification by AutoTT. (D) Cell size and geometry analysis. In (A), n indicates animal numbers. In (B-D), n indicates cell numbers. Student's t-test, *P<0.05, **P<0.01, ***P<0.001, ****P<0.0001. WT, Palmd$^{+/+}$. Het, Palmd$^{+/-}$. KO, Palmd$^{-/-}$.

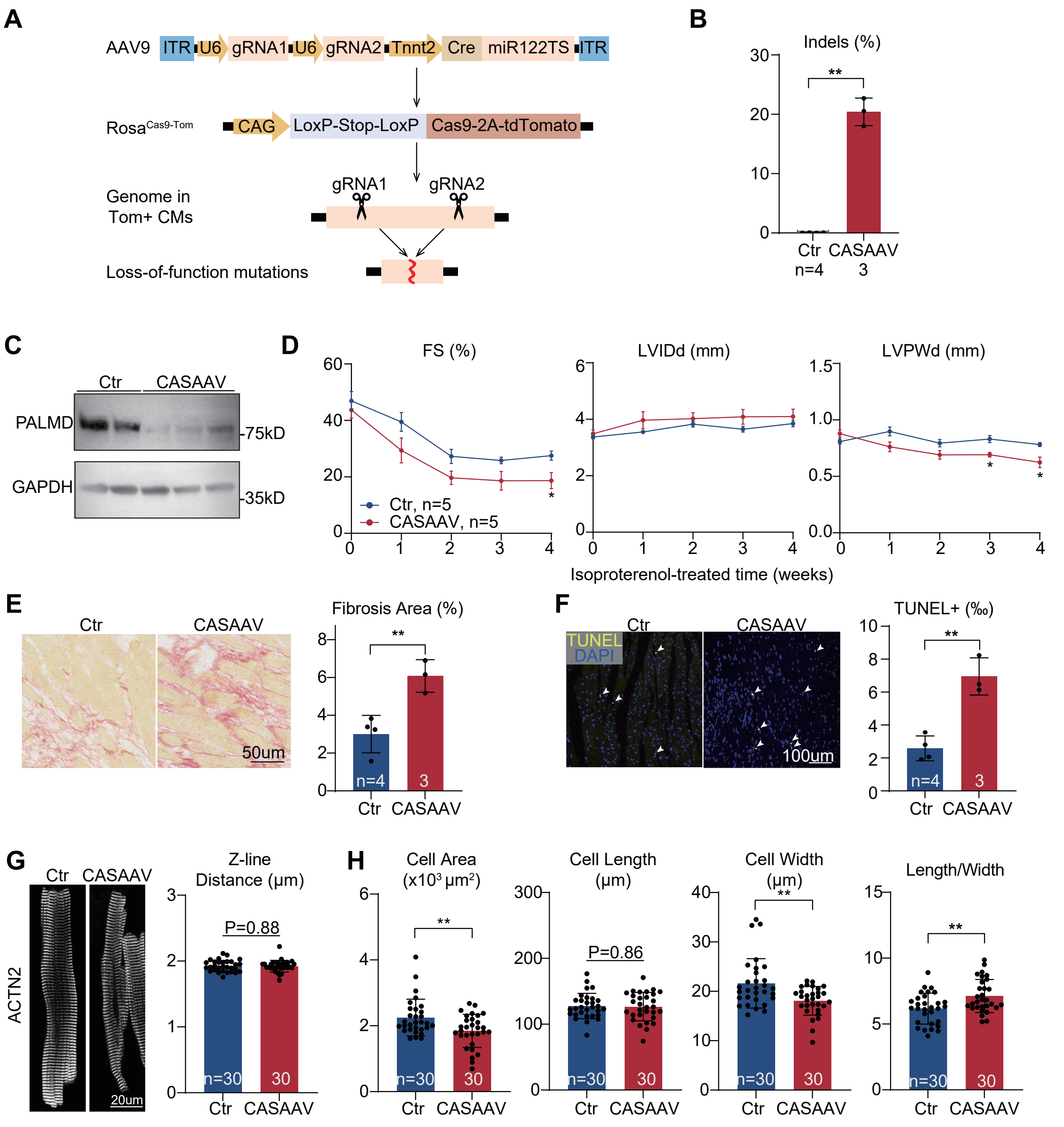


**Figure S6. CASAAV-based cardiomyocyte specific PALMD depletion.** (A) A diagram showing the workflow of CASAAV. (B) Amplicon-sequencing analysis of CASAAV-targeted genome in the heart. Indel, small insertions or deletions. (C) Western blot analysis of CASAAV-treated cardiac tissues. (D) Echocardiogram analysis of CASAAV-treated hearts. (E) Picrosirius red staining and quantification. (F) TUNEL staining and quantification. (G) ACTN2 staining on isolated cardiomyocytes and Z-line distance quantification. (H) Size and geometry analysis of isolated cardiomyocytes. Student's t-test, *P<0.05, **P<0.01, ***P<0.001. In (B, D-F), n indicates animal numbers. In (G-H), n indicates cell numbers. In (D), mean±SEM. In (B, E-H), mean±SD.

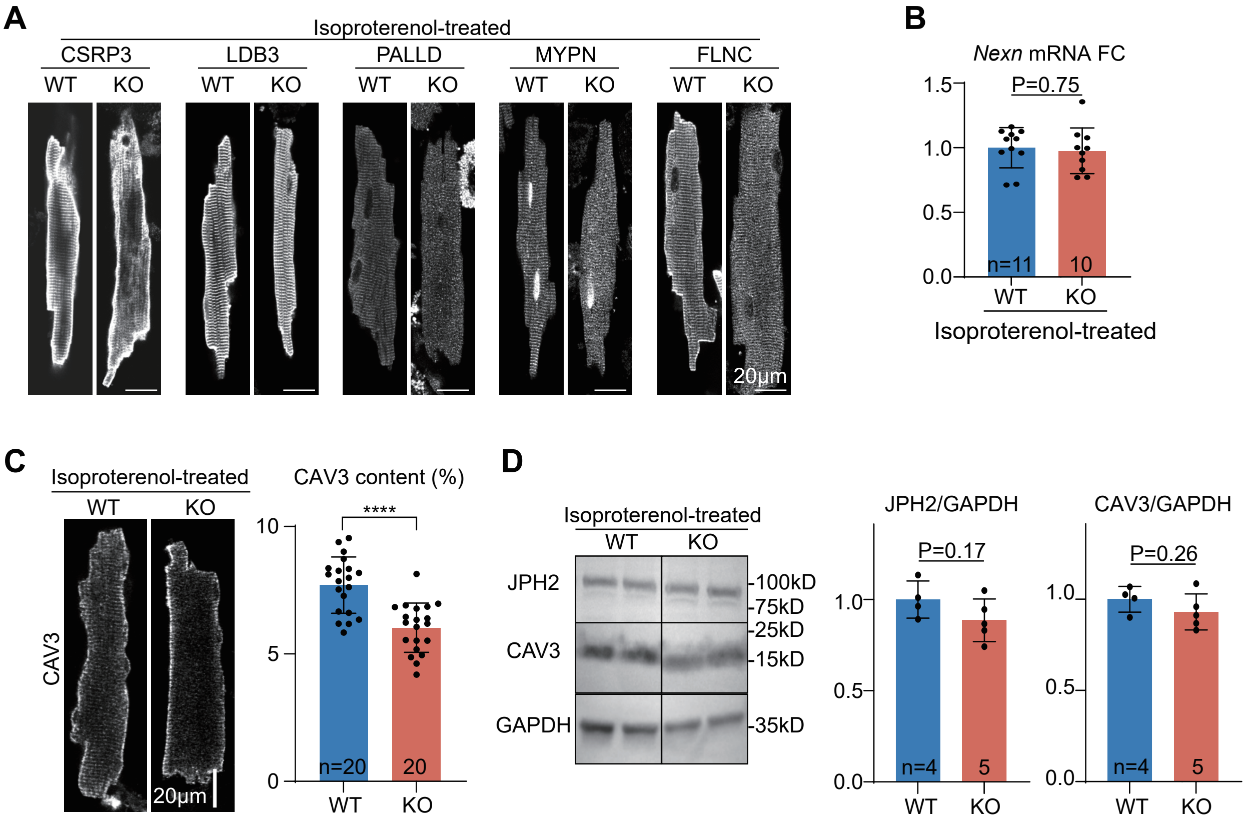


**Figure S7. Sarcomere and T-tubule marker analysis.** (A) Immunostaining analysis of selected Z-line-associated proteins in isolated cardiomyocytes. (B) RT-qPCR analysis of Nexn in hearts. (C) Immunostaining analysis of TT marker CAV3 and AutoTT-based quantification of patterned CAV3 contents. (D) Western blot analysis and quantification. In (B, D), n indicates animal numbers. In (C), n indicates cell numbers. Mean±SD. Student's t-test, ***P<0.001.

### Figure S8

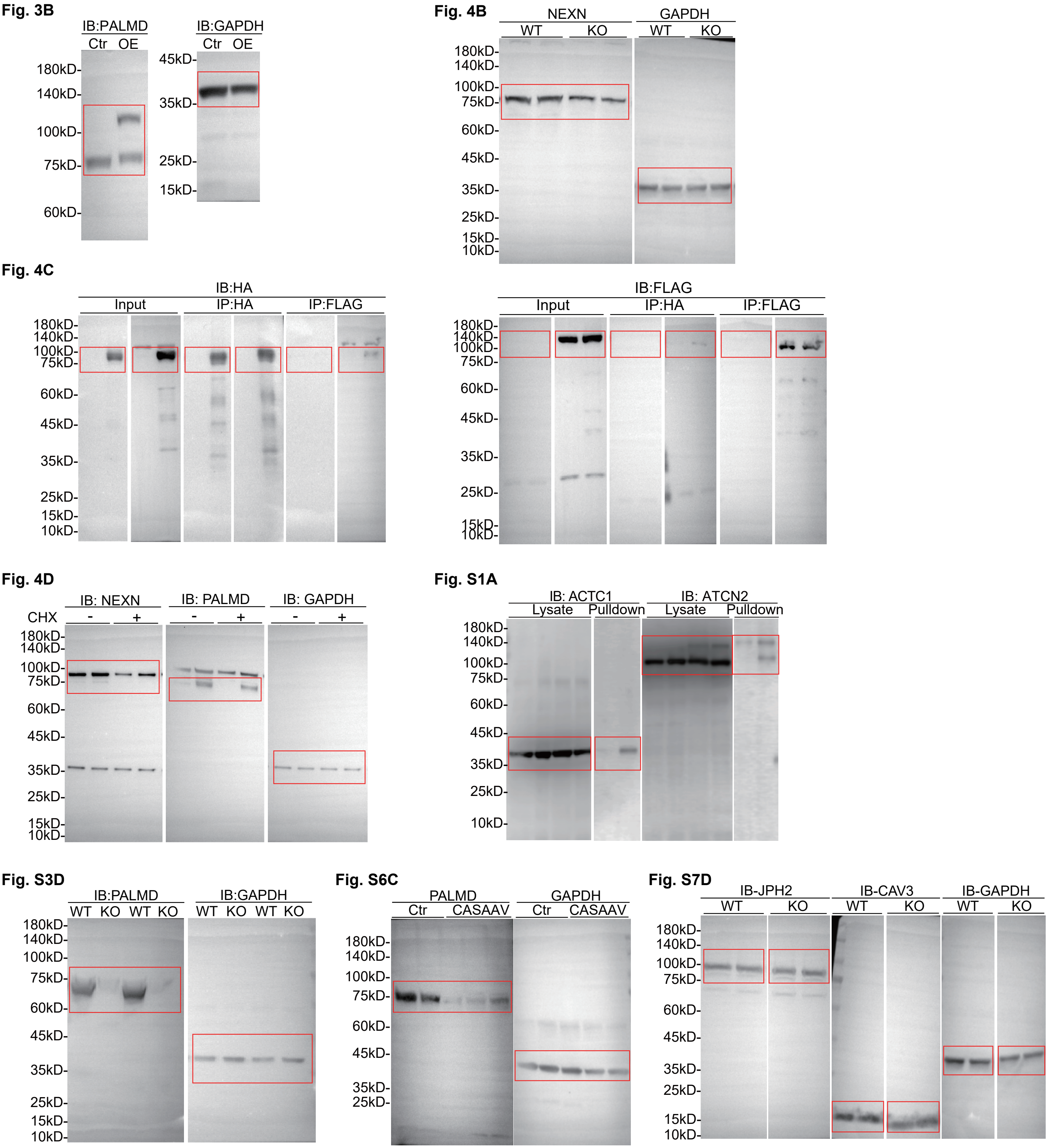


**Figure S8. Full-length raw western blots for each panel.**

**Supplementary Table 1. LC-MS/MS raw data (in a separate file)**

**Supplementary Table 2. Important nucleic acid sequences in this study**

**Genome editing**

| | sgRNA1 | sgRNA2 |
|---|---|---|
| sgRNA in Palmd KO mouse production | 5’-GATGCTTTGCTCTAGAACCT-3’ | 5’-GACATCAGAAAAAAGCCCTG-3’ |
| sgRNA targeting Palmd by CASAAV | 5’- GATGCTTTGCTCTAGAACCT-3’ | 5’-GTTCCACTGCCGATTCCATCC-3’ |

**Palmd Genotyping**

| Genotype | Primer F | Primer R |
|---|---|---|
| WT: 604bp<br>KO: 495bp | 5’-TGGTCTCTGACCTGTGGTCT-3’ | 5’-ACTGAGGAAGACATGCCAGC-3’ |

**RT-qPCR**

| Gene | Primer F | Primer R |
|---|---|---|
| *Palmd* | 5’-CGCCAATCCGTTTTGCAGGC-3’ | 5’-TCCCAAGGGCTTGCCATCCT-3’ |
| *Nexn* | 5’-CAAGCCGGAAATTACATGGT-3’ | 5’-CTCGCTGCTGAGCCTTTATT-3’ |
| *Gapdh* | 5’-TGACCACAGTCCATGCCATC -3’ | 5’-GACGGACACATTGGGGGTAG-3’ |
| *cTNT promoter (* AAV quantification) | 5’-TCGGGATAAAAGCAGTCTGG-3’ | 5’-CCCAAGCTATTGTGTGGCCT-3’ |

**Amplicon sequencing***

| | |
|---|---|
| Barcoded forward primers | 5’-AATGATACGGCGACCACCGAGATCTACAC**TATAGCCT**ACACTCTTTCCCTACACGACGCTCTTCCGATCT**AAAAGTGTATGAGGACCGGC**-3’ |
| | 5’-AATGATACGGCGACCACCGAGATCTACAC**ATAGAGGC**ACACTCTTTCCCTACACGACGCTCTTCCGATCT**AAAAGTGTATGAGGACCGGC**-3’ |
| | 5’-AATGATACGGCGACCACCGAGATCTACAC**CCTATCCT**ACACTCTTTCCCTACACGACGCTCTTCCGATCT**TGTCGCCATGCATAAGACTT**-3’ |
| | 5’-AATGATACGGCGACCACCGAGATCTACAC**GGCTCTGA**ACACTCTTTCCCTACACGACGCTCTTCCGATCT**TGTCGCCATGCATAAGACTT**-3’ |
| Barcoded reverse primers | 5’-CAAGCAGAAGACGGCATACGAGAT**CGAGTAAT**GTGACTGGAGTTCAGACGTGTGCTCTTCCGATCT**CCCAGTCCGTTCGTTTTCAT**-3’ |
| | 5’-CAAGCAGAAGACGGCATACGAGAT**TCTCCGGA**GTGACTGGAGTTCAGACGTGTGCTCTTCCGATCT**CCCAGTCCGTTCGTTTTCAT**-3’ |
| | 5’-CAAGCAGAAGACGGCATACGAGAT**AATGAGCG**GTGACTGGAGTTCAGACGTGTGCTCTTCCGATCT**GAAGGAGCTCGGTTTGAGAA**-3’ |
| | 5’-CAAGCAGAAGACGGCATACGAGAT**GGAATCTC**GTGACTGGAGTTCAGACGTGTGCTCTTCCGATCT**GAAGGAGCTCGGTTTGAGAA**-3’ |

*barcodes and genome-matching sequences in bold

**Supplementary Table 3. Antibodies and Dyes used in this study.**

**Antibodies**

| Name* | Vendor | Cat. # | Working concentration | Application |
|---|---|---|---|---|
| Rb-anti-PALMD | Proteintech | 16531-1-AP | 1:2000/1:200 | WB/IF |
| Ms-anti-ACTN2 | Abcam | ab9465 | 1:2000/1:200 | WB/IF |
| Ms-anti-ACTC1 | Millipore | MABT823 | 1:2500 | WB |
| Rb-anti-CSRP3 | Thermo Scientific | PA529155 | 1:200 | IF |
| Rb-anti-NEXN | Abcam | Ab233267 | 1:2000/1:200 | WB/IF |
| Rb-anti-MYPN | Abmart | PK68590 | 1:100 | IF |
| Rb-anti-PALLD | Proteintech | 10853-1-AP | 1:200 | IF |
| Rb-anti-LDB3 | Proteintech | 11004-1-AP | 1:200 | IF |
| Rb-anti-FLNC | Origene | TA376279 | 1:200 | IF |
| Ms-anti-CAV3 | Abcam | ab173575 | 1:2000/1:200 | WB/IF |
| Rb-anti-JPH2 | Thermo Scientific | 40-5300 | 1:500/1:250 | WB/IF |
| Rb-anti-HA | CellSignalingTechnology | 3724 | 1:2000/1:200 | WB/IF |
| Rb-anti-FLAG | Proteintech | 20543-1-AP | 1:2000 | WB |
| Ms-anti-GAPDH | TransGen | HC301 | 1:5000 | WB |
| HRP-Gt-anti-Ms | TransGen | HS201 | 1:10000 | WB |
| HRP-Gt-anti-Rb | TransGen | HS101-01 | 1:10000 | WB |
| HRP--conjugated Streptavidin | Sangon biotech | D111054 | 1:2000 | WB |
| 488-conjugated Streptavidin | Yeasen | 35103ES60 | 1:500 | IF |
| 488-Dk-anti-Ms | Thermo Scientific | A21202 | 1:500 | IF |
| 555-Dk-anti-Ms | Thermo Scientific | A31570 | 1:500 | IF |

*Rb: rabbit, Ms: mouse, Gt:goat, Dk:donkey

**Dye**

| Name | Vendor | Cat. # | Working concentration | Application |
|---|---|---|---|---|
| FM 4-64 | Thermo Scientific | T3166 | 1ug/ml | in situ imaging |
| Hoechst | Abcam | ab145597 | 10ug/ml | in situ imaging |
| DAPI(1mg/ml) | Thermo Scientific | 62248 | 1:1000 | IF |
| WGA-555(2mg/ml) | AAT Bioquest | 25539 | 1:200 | IF |
| WGA-488(2mg/ml) | AAT Bioquest | 25530 | 1:200 | IF |
